# Overcoming oxygen impermeability in PDMS-free organ-on-a-chip microfluidics with nanoporous thermoplastic

**DOI:** 10.64898/2026.05.22.727128

**Authors:** Franziska Buck, Jeroen Bugter, Gerlinde Kruckenbaum, Insa Staecker, Marina Harzi, Antonina Lavrentieva, Thomas E. Winkler

**Affiliations:** Institute of Microtechnology (IMT), Technische Universität Braunschweig, Braunschweig, Germany; Center of Pharmaceutical Engineering (PVZ), Technische Universität Braunschweig, Braunschweig, Germany; Institute of Technical Chemistry, Leibniz University, 30167 Hannover, Germany; Department of Micro- and Nanosystems (MST), Digital Futures & SciLifeLab, KTH Royal Institute of Technology, Stockholm, Sweden

## Abstract

Oxygen availability is a critical yet all-too-often overlooked variable in organ-on-a-chip (OoC) systems. PDMS-based microfluidics remain the most common approach to facilitating oxygen equilibration with the incubator environment, but the material’s tendency to ad- and absorb small hydrophobic molecules can pose significant concerns for pharmacological and toxicological studies. Yet there remains a lack of alternative gas-exchange materials feasible for OoC integration, even as the use of thermoplastic microfluidics in particular has otherwise proliferated. Here, we present commercially available track-etched nanoporous polycarbonate (50 nm pores, 1.18% porosity, ∼0.1 €/cm^2^) as a practical alternative to polydimethylsiloxane (PDMS) for gas exchange in OoC. We show that nanoporous polycarbonate provides a thermoplastic material with an oxygen permeability of 3290 ± 240 fs mol / kg, over an order of magnitude higher than PDMS. We demonstrate integration into existing lamination-based thermoplastic microfluidic fabrication workflows with sustained leak-free operation well above physiologically relevant pressures. We find that nanoporous polycarbonate does not compromise cell viability, but that high water vapor permeance necessitates a high-humidity environment around the device – though thickness-normalized water vapor permeability is notably similar to PDMS. We validate the OoC application with Caco-2 intestinal epithelial cells by monitoring oxygen levels during the critical cell attachment phase, with nanoporous polycarbonate allowing for maintenance of stable oxygen tension, in stark contrast to severe hypoxia in nonporous controls within 30 minutes. We further show that this uncontrolled hypoxia correlates with a time-delayed increase in cellular hypoxia inducible factor-1 reporter expression. Overall, our findings position nanoporous polycarbonate as a low-cost, mechanically robust, and fabrication-friendly alternative that can bring controlled oxygen availability to PDMS-free microfluidics and OoC.

## Introduction

Oxygen concentration in tissues plays a critical role in regulating a multitude of cellular functions and processes. Whereas the air we breathe is 21% oxygen, oxygen levels in human tissues vary between 1% and 14%,[1], [2] based on the balance between the cells’ nature and inherent metabolic activity and the oxygen supply. The latter is dependent on diffusion in the tissue and (with the exception of airway epithelium) convective blood circulation – with cells being typically located within 100 μm of the nearest capillary, with the distance seldom surpassing 200 μm.[3] Nonphysiological oxygen concentrations – found in vivo with, e.g., stroke or tumors – lead to distinct cellular adaptations, with hypoxia-inducible factor-1 (HIF-1; a ubiquitous transcription factor whose α-subunit degradation is directly regulated by oxygen availability) one of the most well-studied pathways.[4], [5] Despite its significance, oxygen availability is frequently overlooked or not sufficiently accounted for in standard in-vitro studies.[6] Mammalian cells are typically cultured in an effectively hyperoxic environment of approximately 18.9 kPa O_2_ (atmospheric air at 37°C and >90% humidity supplemented with 5% CO_2_). Although incubators with additional oxygen/nitrogen gas control are available to create physioxic or hypoxic environments, they represent a not-insignificant additional cost and infrastructure investment and remain non-standard. It is worth noting also that incubator gas control accounts only for the environment at the media surface of standard culture wells or flasks. To reach cells at the bottom, oxygen must diffuse across several millimeters of media. Especially in dense cultures of metabolically active cells, localized oxygen levels may thus be substantially reduced due to high oxygen consumption.[7], [8] Organ-on-a-chip (OoC) systems have succeeded to better mimic several aspects of in vivo environments; the limited confines of microfluidics, with associated shortened diffusion distances and higher cell-to-medium ratios, open up new possibilities as well as new challenges also for oxygen availability. Relevant examples range from oxic-anoxic interfaces to complex gradients, facilitated by a variety of oxygen delivery and scavenging approaches.[6], [9], [10], [11] At the same time, OoC can take advantage of sensor integration capabilities to provide valuable insight into the true oxygen levels experienced by the cells, as well as their respirative turnover.[10], [12], [13] Together, this has recently led to the realization of closed-loop feedback control.[14]

Still, the main determining factor regarding oxygen availability in OoCs is the material of the microfluidic enclosure. Low-gas-permeable materials such as thermoplastics (as well as the more rarely used glass and silicon) allow OoC to closely approach the in vivo situation in that cellular oxygen availability is somewhat independent of the external atmosphere. As long as the media oxygenation and its perfusion are well-controlled (akin to blood), such systems can achieve physiological oxygen levels with much more compact experimental setups even outside of traditional cell culture incubators.[15] However, fluidic tubing, connections and external ports introduce potential sites for undesired oxygen ingress into the system.[12] Additionally, after cell seeding on a culture surface, stable cell-matrix connections take time to develop.[16], [17] If perfusion is initiated too early, the cells may be washed out of the OoC by the flow of the culture medium before they can properly adhere. Without perfusion, however, the cells may then render the environment overly hypoxic during this critical attachment phase. Therefore, and because incubators (including hypoxia ones) remain more readily available than controlled-oxygenation perfusion setups, highly gas-permeable materials remain perhaps the most prevalent. Specifically, polydimethylsiloxane (PDMS) has been widely and consistently used in OoC fabrication due to its favorable qualities, including besides high gas permeability also biocompatibility, optical transparency, and ease-of-use.[18] It is also an enabling material in many of the more advanced oxygen-managed OoC mentioned above (as an internal gas exchange membrane). One key challenge related to the high permeability, however, is the tendency to ad- and even absorb small hydrophobic molecules, which can include media supplements, endogenous cell signaling molecules, and pharmaceuticals. The magnitude and impact of this strongly depends on the specific experimental design and question,[19] and its effects may be adjusted for computationally, [20] or by surface modification,[21], but this naturally presents a concern for studies of toxicity or therapeutic efficacy.

We thus see a continued need for PDMS-free gas exchange materials in OoC construction. After PDMS and similar silicones, the next most common option are porous fluoropolymers,[23] where chemical or mechanical manipulation can be used to create random, tortuous, high-penetration pore networks in the material. Together with their inherent hydrophobicity, this makes for efficient gas exchange for fluids; however, fluoropolymers continue to place greater demands on microfluidic fabrication and integration than other plastics,[24], [25] and the disordered pore networks lead to high optical scattering and thus poor imaging. Nanoporous (sub-100 nm) membranes from other materials, typically with better-defined vertically-aligned pores, provide an alternative that has received the least attention to date.[26] Anodized aluminum oxide was demonstrated to enable high gas permeability in PDMS-free OoC devices, but required multi-step cleanroom-based processing, remains fragile (brittle), and lacked experimental validation (beyond cell viability).[27] Silicon nitride membranes were shown to be superior to PDMS membranes for use in microfluidic blood-oxygen exchangers, but suffer the same material limitations as aluminum oxide.[28] For the same application, another group had previously evaluated nanoporous polycarbonate, which can be mass-produced relatively cheaply, and as a plastic is more resilient. The membranes did not perform well for the authors’ application, however, remaining inferior to both regular and porous PDMS (which their study was focused on).[29]

Here, we set out to establish that nanoporous polycarbonate (which we term NanoPC) is well-suited to facilitate oxygen exchange in PDMS-free OoC. We first establish the material’s oxygen permeability (gas/gas) both experimentally and theoretically to facilitate comparisons with the literature. We proceed to demonstrate NanoPC integration into microfluidics by means of lamination using tape or off-stoichiometry thiol-ene-epoxy. We validate that devices maintain structural and fluidic integrity under relevant pressures, and that liquid/gas exchange remains highly efficient in such devices. After establishing NanoPC biocompatibility and further investigating water vapor permeability, we demonstrate culture of Caco-2 intestinal epithelial cells in corresponding OoC. By continuously measuring cell culture medium dissolved oxygen partial pressure (i.e., oxygen tension), and tracking fluorescent expression of a hypoxia reporter element, we establish NanoPC’s potential for PDMS-free, incubator-equilibrated OoC studies.

## Materials & Methods

### Membranes

Track-etched polycarbonate membranes, 25 µm thick with a nominal 6×10^8^/cm^2^ pores of 50 nm diameter, lacking post-process hydrophilic treatment, were purchased from it4ip (Belgium), termed NanoPC throughout. We illustrate the membrane in Fig. 1 by means of an atomic force micrograph, conducted using a NaniteAFM (Nanosurf, Switzerland) in dynamic force mode using TapAl190-G cantilevers (BudgetSensors, Bulgaria), with data post-processed (level and contrast corrections) in Gwyddion. Native, nonporous polycarbonate (125 µm thick; Makrofol DE 1-1) serves as a control material throughout our study and was kindly supplied by Covestro (Germany).

**Figure 1.**
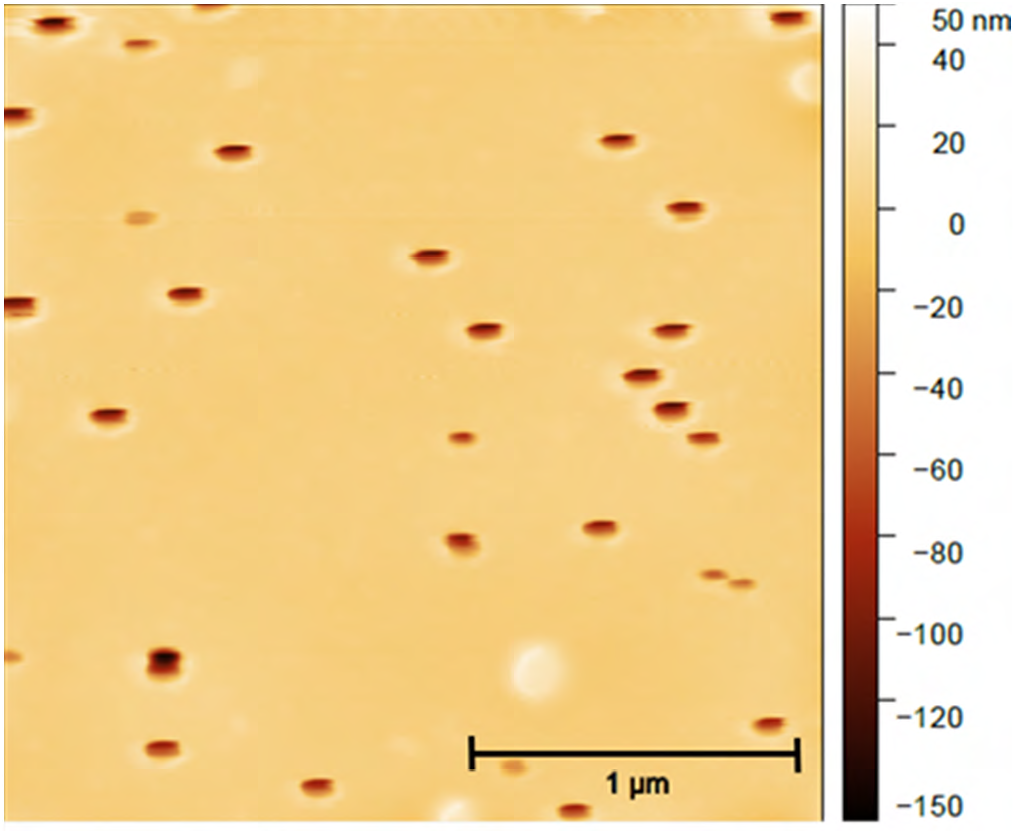
Representative atomic force micrograph of NanoPC, illustrating randomly-distributed pores in the surface in line with the nominal material specifications. The oval appearance of pores is an artifact of the difference in fast-scan and slow-scan axes.

### Sensors

For oxygen measurements, we rely on phosphorescent sensor spots (OXSP5; Pyroscience, Germany) with fiber-optic readout (Firesting FSO2-C4, Pyroscience, Germany), and custom 3D-printed rigs to keep fibers in place. The oxygen readouts were calibrated according to manufacturer instructions and recorded using their proprietary software. Upper-level calibration specifically involved a sealed vial filled with atmospheric air and water, heated up to 37°C inside an incubator, and shaken for 1 minute to fully aerate the water (equaling a theoretical 22 kPa dissolved oxygen partial pressure) – a procedure that involves a tradeoff between reaching full aeration (longer shaking) and limiting room-temperature exposure causing temperature mismatch (shorter shaking).

### Cells

To illustrate cell culture compatibility, we relied on the widely-used intestinal epithelial line Caco-2 as a model (after disinfection and collagen I coating). Specifically, we used C2BBe1, a clonal line, provided by ATCC (USA); for microscopy validation of hypoxia, we additionally employed Caco-2 transduced with HRE-dUnaG,[30] a fluorescent hypoxia reporting element, using a protocol described previously.[31] Further details on cell culture can be found in the SI.

### Test Devices and OoC

NanoPC (and PC controls) were evaluated and/or integrated in a variety of test systems and devices. For initial oxygen permeability evaluation, we mounted 5.5 cm diameter NanoPC on top of a sensor spot-integrated, nitrogen-flushable test cell, illustrated in Fig. S1. For cytotoxicity testing, hanging inserts for 24-well tissue culture plates (Sarstedt, Germany) were modified by excising existing membranes and instead mounting NanoPC or PC samples with ∼6 mm diameter using medical-grade cyanoacrylate adhesive (Cyberbond 2008; H.B.Fuller, USA). For water evaporation tests, NanoPC or PC (or PDMS) were mounted on top of custom 3D-printed vessels with ∼2 ml volume (further details in SI).

All other experiments were conducted using custom-made single-chamber/channel microfluidic devices where NanoPC served as either bottom (pressure and permeability tests) or ceiling (oxygen depletion during cell culture). The various design cross-sections, based on the varying experimental requirements, are illustrated in Fig. 2. All were based on xurography- and lamination-based fabrication with medical tape (Medical Tape 9877; 3M, USA), as described previously;[32] for burst pressure, we additionally consider off-stoichiometry thiol-ene-epoxy (OSTEmer 322; Mercene, Sweden)-based fabrication to illustrate the extensibility of NanoPC integration.[33] Additional fabrication details can be found in the SI. Devices with pressure or microscopy endpoints (a,d) featured the simplest designs; those that called for continuous oxygen measurement employed additional layers to facilitate sensor spot integration. Excepting the pressure test devices (i.e., a), microfluidic channels and chambers of interest had a height of 1 tape layer, equating approximately 110 µm.

**Figure 2.**
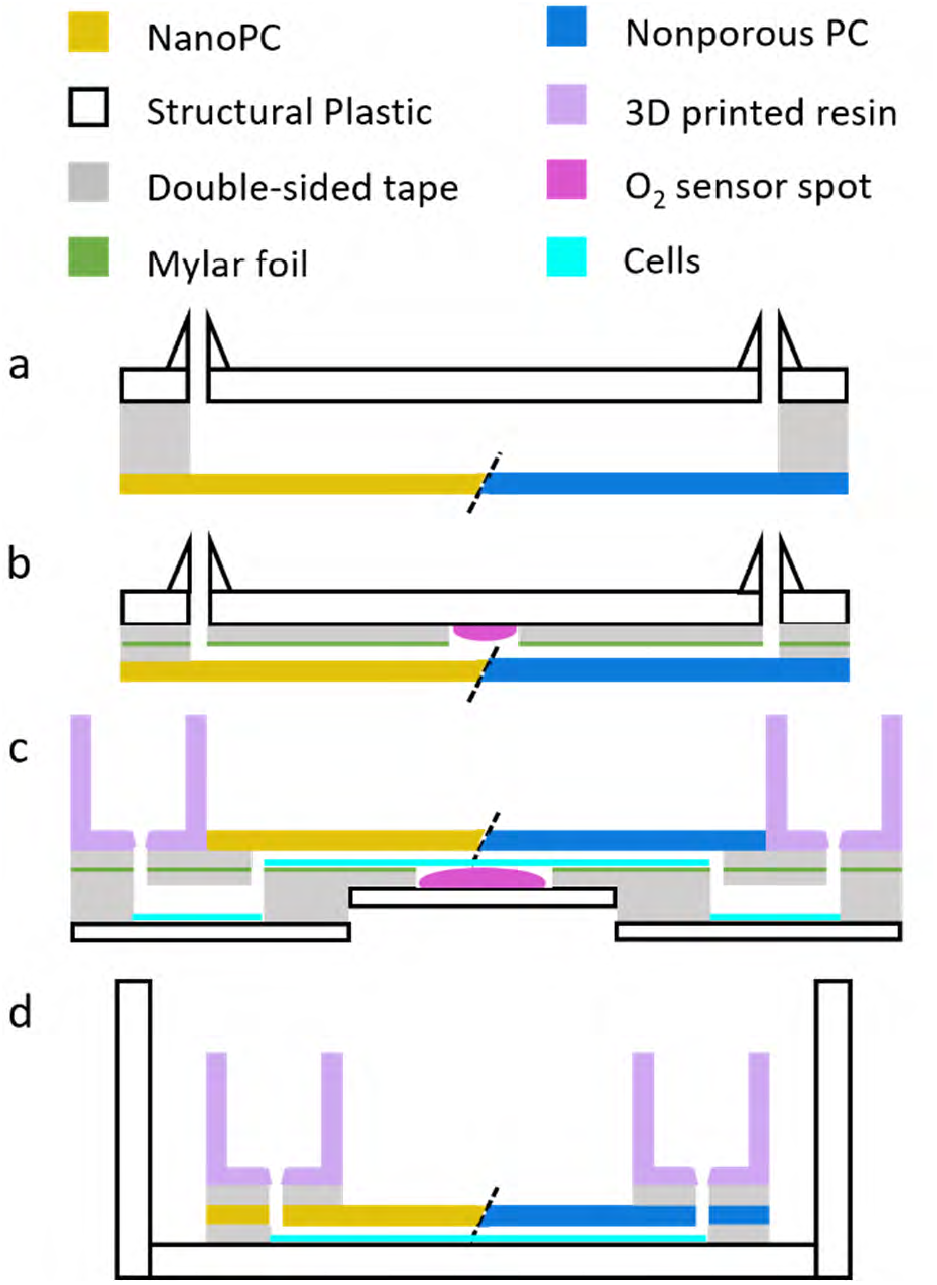
Length-wise cross-sections of the microfluidic devices and architectures used in our study.

### Quantitative Analysis

Quantitative data were analyzed and visualized in OriginLab Pro unless otherwise noted. Any p-values and associated statistical significance were evaluated using one-way ANOVA with Tukey post-hoc means comparison.

### NanoPC membranes: oxygen permeability, water vapor permeability, cytotoxicity

Theoretical permeability calculations assumed a combination of bulk and Knudsen diffusion through cylindrical pores, and is detailed in the SI. Experimental test cells were prepared with a thin layer of silicone-free grease, NanoPC samples laid on top, and any trapped air removed using a scraper. After cell closure, the setup was flushed with N_2_ for 5 min, followed by 3 min of measurement. Corresponding data analysis is described in Fig. S1. Membrane oxygen permeability tests are presented in terms of arithmetic mean and standard deviation to provide consistency with literature values.

Water permeance and permeability were assessed by filling the 3D-printed reservoirs with water and mounting membranes on top, sealed by an O-ring. Reservoirs were weighed, placed in a climate-controlled room (22°C, 23–28% humidity), and re-weighed after 24 h. In calculating permeance and permeability from the resulting rates, we account for open-reservoir controls by treating the membrane as a series resistance.

For cytotoxicity evaluation, Caco-2 were seeded onto the well plate inserts at a density of 15×10^6^/cm^2^. After ∼24 hours, cells were stained with Calcein-AM (TOCRIS Bioscience, UK) and Propidium Iodide (Avantor, USA). Optical images were taken using an Olympus IX73 inverted microscope and post-processed in Fiji/ImageJ. Due to variable live/dead staining efficiency, image brightness and contrast were linearly adjusted per-image to yield perceptually similar contrast across the dataset.

### NanoPC device integration: pulse pressure, sustained pressure, and oxygen permeability

Devices shown in Fig. 2a were filled with deionized water (mixed with food coloring). Outlets were clamped shut, and inlets were connected to a microfluidic flow controller and in-line flow sensor (Flow EZ or MFCS EZ, and Flow Unit M; Fluigent, France). The proprietary instrument software was used to set target pressures, and to record flow. For pulse tests, pressure on the devices was increased in a stepwise fashion (10 kPa per 10 s) until 300 kPa (tape devices) or membrane rupture (OSTEmer devices). Failure points were defined as the pressure where flow exceeded the baseline (employing a 1.96σ cutoff criterion for tape-based devices; Fig. S2). For sustained tests, pressure was set to either 20, 30, or 40 kPa over 24 hours. Leakage was determined visually, either by seeing drops leaking from the system, or by colored stains left on tissue paper under the devices. We fit our failure probability distribution with a normal cumulative distribution function to derive the 1.96σ-analogous 2.5% failure pressure (Fig. S3).

Devices shown in Fig. 2b were filled with phosphate-buffered saline (PBS; pre-incubated in a standard cell culture incubator), and placed in a nitrogen-flooded cell culture incubator, where oxygen data was recorded over 24 hours. Dissolved oxygen partial pressure time series data were processed in RStudio as follows: log-space transform (to account for first-order exponential nature of data); 60 s moving-median with 30 s downsampling (to account for transient noise); two-stage (sample-replicate) bootstrap calculation of (geometric) means, and calculation of median and interquartile ranges (IQR) on resulting population (thus capturing both between-sample and within-sample variability). Outliers were included in the analysis, but were identified as such if a certain fraction of data (corresponding to the characteristic time constant of the shortest median exponential decay of a given experiment) lay outside the Tukey fence.

### NanoPC-OoC: oxygen sensing and microscopic observation

Given the opaque nature of the oxygen sensors, we chose to employ two distinct device designs optimized for either media oxygen monitoring (Fig. 2c) or cell hypoxia observation (Fig. 2d). Cells were seeded at a density of 15×10^6^/cm^2^, and recordings started with an estimated delay of 5 minutes. Oxygen data were recorded for 6 h; in corresponding devices, presence of cells was validated by brightfield observation in between inlets/outlets and sensor spot. Microscopic observations for hypoxia were conducted at approximately 6 h and 26 h using an Olympus IX73 inverted microscope and post-processed in Fiji/ImageJ, adjusting brightness and contrast linearly on a per-experiment basis. All solutions were pre-incubated for at least 1 h prior to use, and great care was taken to ensure a high-humidity environment in the immediate OoC vicinity by filling adjacent wells and reservoirs with water.

## Results & Discussion

Polycarbonate (PC) is a widely-utilized thermoplastic, both generally as well as for microfluidic engineering. In OoC, it is encountered not only in the microfluidic enclosure, but also as a permeable cell culture support, serving to support cell adhesion and to provide compartmentalization.[34] These permeable supports are created by chemically etching pores (often in the range of 0.4 to 3 µm diameter) along the vertical tracks left by high-energy ion bombardment (i.e., ion track etching) through a ≲30 µm film, and subsequently applying a hydrophilic cell culture surface treatment.[35] The same high-throughput process can however also yield membranes with pores in the tens of nanometers, and without surface treatment, and such membranes are similarly available commercially at low cost (∼€0.1/cm^2^).

We hypothesize that this NanoPC can combine a number of desirable properties for fabricating PDMS-free, gas-permeable OoC:

1. it inherits the beneficial properties of widely-used PC and PC-based permeable culture supports, including but not limited to: range of bonding and fabrication strategies, favorable mechanic properties for handling and resilience, optical clarity, biocompatibility, autoclavability, and low cost;
2. physical pores providing oxygen permeability (we select *ε*=1.18% porosity);
3. zero microbial permeability based on pore size (*d*=50 nm);
4. no optical degradation due to well-ordered sub-wavelength pores; and
5. low water permeability based on high aspect ratio pores (*t*=25 µm thickness, thus 1:500) with near-90° surface contact angle (higher than that of, e.g., track-etched PET).

### NanoPC offers increased oxygen permeability over PC and PDMS

We first seek to establish the fundamental oxygen permeability *P*_O2_ of NanoPC and to place this in the context of alternative material options. For NanoPC, we pursue both a theoretical estimation based on gaseous diffusion through cylindrical pores, and an experimental evaluation. For the latter, we relied on a custom-built test cell with the membrane clamped between ambient air on one side, and a gas- and flow-controlled chamber with an oxygen sensor on the other, allowing us to extract permeability from dynamic oxygen equilibration data. Data for alternative materials were extracted from the literature, converted as appropriate, and confounding factors noted. The results are shown in Table 1.

**Table 1.** Oxygen permeability of NanoPC (this study) as well as literature values for other relevant materials (for selection rationale, see text). NanoPC experimental readings are mean ± SD from 3 measurements each on 3 samples; we elaborate on the theoretical calculation in the SI. We translated literature values into equivalent units to facilitate comparison. Temperatures are denoted as this can impact permeability (for bulk materials, we generally expect higher permeability at higher temperature), with ? indicating lacking specification. We moreover denote additional properties of the materials with relevance for use as external microfluidic enclosures.

| Material | Material notes | $P_{O_2}$<br>[$10^{-15} \text{ s mol / kg}$] | $T$ [°C] | Experimental notes |
| --- | --- | --- | --- | --- |
| NanoPC (this work) | Plastic<br>$\sim 90^\circ$ contact<br>angle | 26 140 | 23 | Theory |
|  |  | 25 940 | 37 | Theory |
|  |  | **3 290 ± 240** | room | Experiment |
| Polycarbonate (PC) |  | 0.46 | 25 | Lexan [37] |
|  |  | 0.33 | 37 | Lexan [38] |
| Polydimethylsiloxane (PDMS) | Small-molecule<br>sorption | 180 ± 20 | room | Sylgard 184 [39] |
|  |  | 201 | 40 | RTV 615 [40] |
|  |  | 245 | 37 | Elastosil 2030 250 [38] |
| Styrene-ethylene-butylene-styrene (SEBS) | Elastic | 3.10 | 23 | Kraton G1643 [41] |
|  |  | 3.21 | 23 | Kraton G1645 [41] |
|  |  | 7.86 | ? | FlexDym [42] |
| Polymethylpentene (PMP) | Elastic | 10.4 | 37 | TPX DX845 [38] |
| Polyurethane (PU) | Elastic | 0.20–0.63 | 25 | PBA-based; various ratios [43] |
|  |  | 0.39–3.53 | 25 | PTMG-based; various ratios [43] |
|  |  | 9.04 | 37 | PPO-based [44] |
| Porous fluoropolymer | Plastic | 26 500 ± 1 000 | ? | PVDF [45] |
|  | Heterogeneous pores | 812 000 ± 19 000 | ? | HFP5 [46] |
| Microporous silicon nitride | Brittle<br>Very hydrophilic<br>Non-sterile | 152 000 | ? | Unclear if theory or experimental [28] |
| Nanoporous anodized aluminum oxide | Brittle<br>Very hydrophilic | 316 000 | 37 | Theoretical estimate [27] |

We thus find that NanoPC is orders of magnitude more oxygen permeable than solid PC, confirming the fundamental principle. Furthermore, our measurements place NanoPC at over an order of magnitude more oxygen permeable than solid PDMS. We emphasize that permeability is thickness-normalized, and that silicone membranes are more prone to tearing at equivalent thickness (∼10-fold lower tensile strength;[36] i.e., as an external enclosure material, they would have to be thicker for equivalent device resilience). Our result presents a curious yet clear contrast to that of Wu et al., who employed thinner (6 µm) but otherwise nominally identical NanoPC in a blood-oxygen exchanger.[29] Although they do not directly assess *P*_O2_, they found that the blood-oxygen exchange rate across NanoPC was at best not significantly different from, and at worst a significant 40% lower than, 15 µm thick PDMS. The fact that another material (closed-pore PDMS) showed superior performance rules out measurement-related saturation effects. Part of the discrepancy may be due to the specific NanoPC membrane used (i.e., deviations from nominal), as well as their membrane integration process relying on uncured PDMS (applied at bonding points only, but with potential to spread and clog adjacent pores). We hypothesize that the specific application does explain the largest part, however, as blood is a much more complex fluid than any of the ones we consider herein, containing in particular many more components that may lead to pore clogging (e.g., coagulation factors, platelets, etc.).

Although we find NanoPC to be highly oxygen permeable both experimentally and theoretically, the experimental value lags behind the model prediction by a factor of ∼8. A small fraction of this may be due to fundamental assumptions and simplifications in our model (e.g., neglecting entrance effects). Moreover, track-etched pores are not perfectly cylindrical, and expected nanometer-scale deviations towards (bi)conical profiles will have sizable impact on transport properties at our chosen pore diameters. As with the uncured-PDMS bonding mentioned above, grease or oil from the experimental setup and manual handling are also likely to contaminate a fraction of the pores. What we judge most significant, though, is that residual back-pressure from the nitrogen flow prior to the measurement is likely to slow down measured diffusion.

Beyond NanoPC and PDMS, we further illustrate the poor *P*_O2_ of materials considered as PDMS alternatives for microfluidic fabrication in Table 1. SEBS, PMP, and PU have all been investigated and to varying degrees found acceptance, given that they offer superior elasticity and oxygen permeability to thermoplastics like PC, and reduced small-molecule sorption to PDMS. Yet they still fall far short of PDMS (and NanoPC) on *P*_O2_. Regarding PDMS, we also note here that a myriad strategies for its surface modification have been proposed, which can mitigate small-molecule sorption;[21], [22] some of these are known to in turn reduce *P*_O2_, but for most this has not been evaluated to date. The longevity, robustness, and added effort needed to realize such coatings however continue to pose limitations.

Porous fluoropolymers can conversely achieve very high *P*_O2_ due to their very high pore density. As mentioned earlier, they suffer from poor transparency and being less amenable to microfluidic fabrication process integration [24], [25]. Ordered pores can address the former, especially when coupled to thinner membranes. One notable example is that by Imtiaz et al., who illustrate excellent performance of 400 nm thick silicon nitride in the context of blood-oxygen exchange.[28] They however operate the membrane as a controlled *liquid-liquid* interface – given that their 0.5 µm pore size (along with inherent hydrophilicity) cannot prevent liquid leakage nor microbial ingress. While smaller pores are also possible in the same cleanroom-based fabrication process (and similarly commercially available), the associated high cost of ∼€50 per membrane with an effective surface area in the mm^2^-range stands in stark contrast to all other materials. Bunge et al., meanwhile, consider 1.5–3 µm thick anodized aluminum oxide featuring 38 nm pores in the context of OoC.[27] This is sufficient to maintain sterility, and although the hydrophilic pores still wet, they do maintain liquid integrity. The authors’ characterization is limited to showing cell viability, but we can estimate at least a theoretical *P*_O2_ that is well above NanoPC. Their in-house fabrication process is, however, also cleanroom-based and resource-intensive. The commercial pricing of ∼€10/cm^2^-scale membrane is certainly an improvement over silicon nitride, but still far short of €0.1/cm^2^ for track-etched thermoplastic. Finally, both silicon nitride and aluminum oxide membranes as described in the literature are exceedingly fragile due to their thinness and brittleness, and thus not suited for use as external microfluidic materials.

### NanoPC-integrated OoC maintain liquid integrity under typical operation pressures

Having established that track-etched nanopores, i.e., NanoPC, are a promising approach to facilitate gas permeability in thermoplastics, we proceed to evaluate its potential for microfluidic and OoC device integration. We do this by assessing how NanoPC affects the device leakage compared to native PC, both (1) short-term, under rapid pressure bursts (i.e., as may be employed to flush a microfluidic device), and (2) long-term, under sustained 24 h liquid pressure (i.e., mimicking continuous perfusion). We specifically consider two methods of plastics-based microfluidic fabrication amenable to NanoPC integration:

a. layer-by-layer lamination of PC and precision-cut double-sided tape,[32] one of the most cost-efficient approaches for thermoplastic OoC fabrication and assembly; and
b. reaction-injection-molding of OSTEmer,[33] a facile polymer for OoC fabrication, given its excellent inherent multi-material thermal bonding capabilities [47] and low-sorption end-state [48]; it is however also gas-impermeable (and can be made to even scavenge oxygen).[47]

For burst pressure testing, we rely on flow sensor readings under increasing dead-end hydrostatic pressure to determine the point of microfluidic device failure. Fig. 3a illustrates that, in otherwise identical OSTEmer devices, NanoPC performs significantly worse than PC, but significantly above 100 kPa. For tape devices with equivalent geometry, the burst pressure remains around 100 kPa in both cases, without statistically-significant differences between NanoPC and PC. This indicates a burst pressure failure inherent to the tape itself, while the differential results with OSTEmer point to a NanoPC-inherent failure. Indeed, both from the sensor recordings (Fig. S2) and visual inspection, we observe that tape devices fail by gradual delamination of the PC or NanoPC, in contrast to OSTEmer devices that fail by NanoPC membrane rupture.

**Figure 3.**
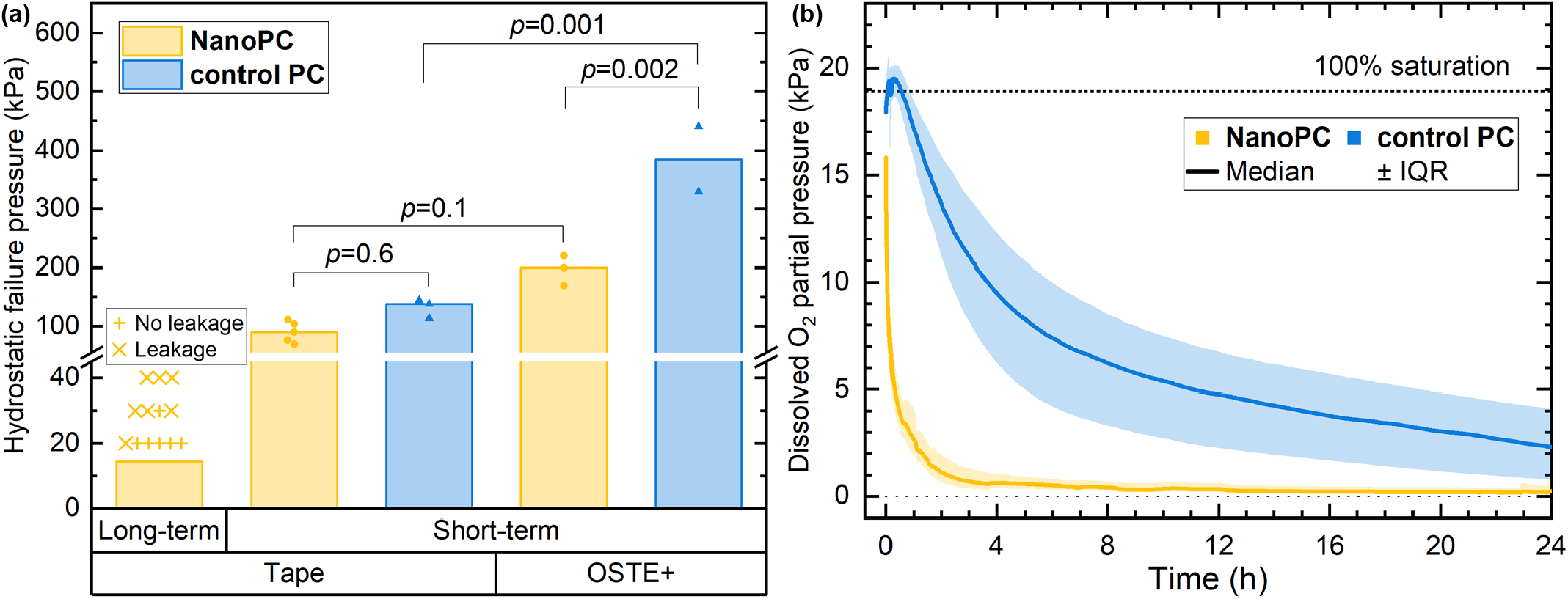
NanoPC microfluidic device integration tests. (a) Pressure resilience using tape- or OSTEmer-based device fabrication, under either short-term pulse pressure or sustained pressure, comparing NanoPC against PC controls. For short-term pulse tests, each point represents a single device failure pressure (bars at median level), with p-values derived from ANOVA post-hoc Tukey test. For sustained pressure, every + indicates a single device that did not leak after 24h, and every x represents a single device that did. The bar represents the estimated 2.5% failure pressure (Fig. S3). (b) Oxygen tension recorded inside NanoPC- and control PC-integrated tape microfluidics after placement inside a nitrogen-flooded cell culture incubator, illustrating the orders-of-magnitude different equilibration timescales. The line represents the median of bootstrap means across experiments (NanoPC: 18; control PC: 7), with shaded regions indicating corresponding IQR.

For sustained pressure evaluation, the leakage flow proved too low to be measured by our flow sensor. Instead, we rely on a simple yes/no classification based on whether we observe residue from the food color-supplemented water on a tissue placed underneath (Fig. S2). From the failure probabilities at 20, 30, or 40 kPa over 24 h we can estimate an onset failure pressure (P2.5) of 13.3 kPa (Fig. S3). Put into perspective, this is 2.5-fold higher than the typical blood pulse pressure, and is sufficient to achieve a wall shear stress of 33 dyn/cm^2^ over a 1 cm long channel with a diameter as low as 10 µm. The higher short-term burst pressure resilience moreover supports set-up and maintenance operations like bubble flushing. We thus conclude that NanoPC is well-suited for OoC device integration.

We also expect that the evaluated fabrication methods can be further extended to other thermoplastic assembly methods.[25] Solvent bonding is likely not advisable; however, controlled adhesive bonding (being very similar to the tape approach) and ultrasonic bonding are expected to be suitable. Even thermal fusion bonding, one of the most attractive methods for academic thermoplastic bonding due to its simplicity, can be adapted. However, elevating the temperature above the glass transition temperature *T*_g_ will cause the pores to shrink due to the Laplace pressure exceeding the zero-shear viscosity. Thus, the temperature differential compared to *T*_g_ should be kept small, and a larger initial membrane pore size should be chosen to compensate for remaining shrinkage.

### NanoPC-integrated OoC maintain high oxygen permeability

To further establish that the oxygen permeability of NanoPC functions as intended for OoC application, we measure the oxygen tension over time using fully-assembled (tape-based) microfluidic devices, now integrated with an optical oxygen sensor. In Fig. 3b, we display oxygen recordings as devices filled with PBS (pre-incubated to atmospheric air, 5% CO_2_, 37°C, >90% humidity) were moved to an otherwise equivalent hypoxia incubator environment (0% O_2_). The orders-of-magnitude difference in equilibration time constant – 10.1 minutes for NanoPC, 6.5 hours for native PC, assessed relative to the theoretically-expected limit of 18.9 kPa from the standard incubator environment – confirms that high NanoPC permeability is maintained with OoC integration and application. The native PC, meanwhile, is clearly limited by the long-distance (>1 cm) and low-area (<1 mm^2^) oxygen diffusion from the open channel inlets to the centrally-placed oxygen sensor.

For the native PC, we also note an initial overshoot in the oxygen measurement, going above the 18.9 kPa “limit”. We attribute this to imperfect sensor calibration, given that the procedure inadvertently introduces small errors in both temperature (1–2°C below nominal 37°C) and oxygen tension (few percent below limit; cf. Methods). The test devices – with the small internal solution volume cooling down between device loading and incubator introduction – are thus expected to start out at or below the true calibration conditions, matching observations. As both devices and incubator (also subject to thermal loss from door-opening) thermally re-equilibrate, oxygen readings initially increase as true 37°C is approached. This thermal process occurs in parallel with gas equilibration towards nitrogen atmosphere, which with NanoPC is rapid enough to obscure this initial artifact.

### NanoPC does not affect cell viability, but water evaporation does

While PC is generally considered biocompatible, the NanoPC we employ has not previously been evaluated for cell culture. We thus consider its impact on cell viability first by establishing a Transwell-style system where the permeable culture support was replaced by either NanoPC or native PC. In Fig. 4a, we illustrate representative live/dead staining of Caco-2 cells on these materials. We find no observable differences in cell coverage or viability between the two materials when media is present in both the inner and outer compartments. However, when exposed to air underneath, only cells on solid PC maintained high viability, whereas cells on NanoPC exhibited wide-spread cell death. We attribute this to water vapor permeability of the NanoPC, which would render the cells hyperosmotic.

**Figure 4.**
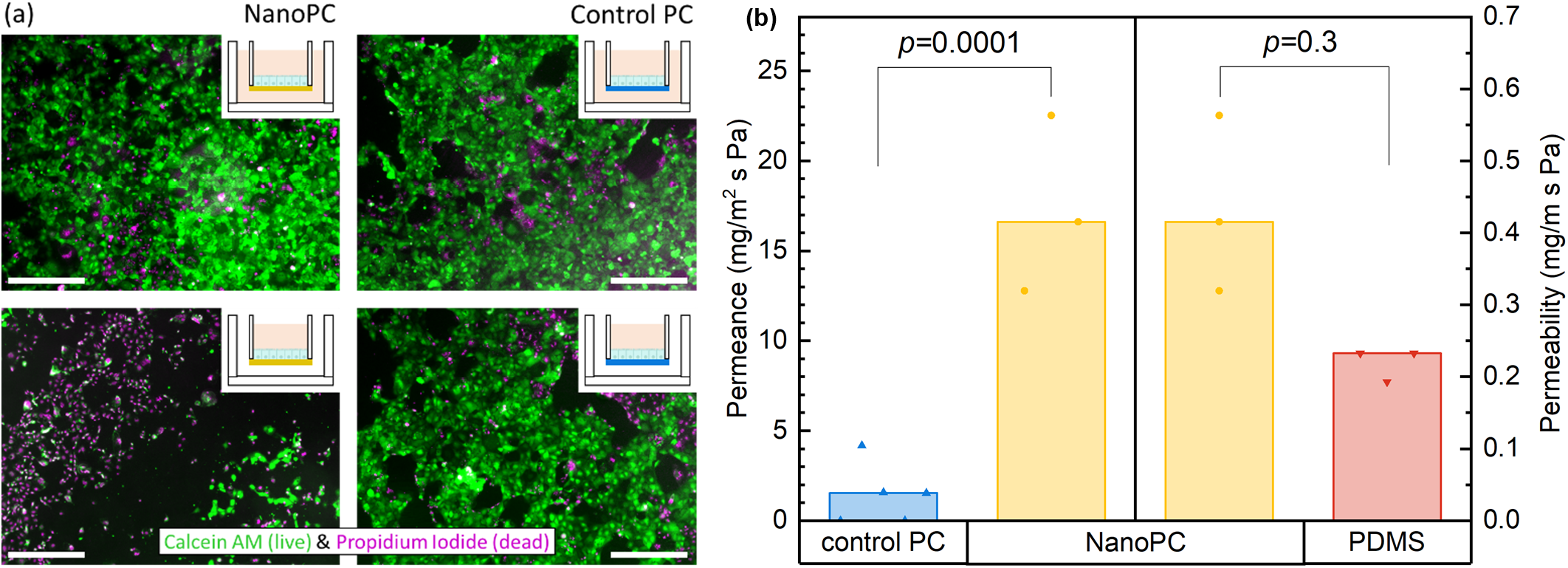
(a) Cytotoxicity evaluation of NanoPC (left) as compared to PC controls (right) employed as cell culture substrates in Transwell-like geometry. Caco-2 cells are fluorescently stained with a membrane-permeant dye-linked esterase substrate (green; live cells), and a membrane-impermeable DNA-intercalating dye (magenta; dead cells). Representative micrographs reveal no inherent difference between NanoPC and control when immersed in liquid (top); with air underneath (bottom), NanoPC performs significantly worse, suggesting a potential water-evaporation effect. Scale bars: 500 µm. Additional images in Fig. S4. (b) Water vapor permeance for NanoPC and PC controls, as well as thickness-normalized permeability for NanoPC and PDMS controls. Each dot represents a membrane sample (bars at median level), with p-values derived from ANOVA post-hoc Tukey test. This confirms significantly increased water loss through NanoPC.

Indeed, we find that the water vapor permeance through NanoPC is 11-fold as great as that of native PC, in line with our hypothesis (Fig. 4b). At the same time, however, we notably find no significant difference between NanoPC and PDMS on a thickness-normalized basis (i.e., permeability). We emphasize that this is in great contrast to the substantial differences in gas permeability *P*_O2_, confirming our assumption about the hydrophobic pores helping to mitigate water transport. Still, the absolute increase in water vapor permeance is sufficient to impact cell viability. While direct impact on cells can be mitigated by simply not culturing them directly on the NanoPC but instead on a distal surface, on a permeable support, or distributed in a 3D hydrogel (all relevant for OoC), this still retains a risk of detrimental hyperosmotic conditions (with volume-limited, static media). It thus becomes important to ensure a high-humidity environment around the NanoPC, which – due to the de-coupled recovery dynamics of temperature and humidity across the incubator volume – cannot be ensured by relying on the incubator alone. We address this by placing liquid-filled reservoirs in device proximity, and adding moist tissues or water drops on the devices themselves in subsequent experiments.

Curiously, Bunge et al. did not observe wide-spread cell death on their anodized aluminum oxide membranes (though their experiment lacks a control condition).[27] Yet their membrane should suffer from even higher evaporative loss (∼20-fold higher porosity, fully-wetted pores). We expect this can in part be attributed to their choice of immortalized keratinocytes (HaCaT), which are known to thrive under skin-appropriate air-liquid interface culture. Moreover, their OoC featured a confined air channel, which would only require marginal media evaporation to re-equilibrate after temperature fluctuation, providing a buffer with regard to the larger (slower-equilibrating) incubator environment.

### NanoPC-integrated OoC ensure incubator-equilibrated culture conditions and prevent excess hypoxia in the attachment phase

Ultimately, we proceed to demonstrate the OoC application of NanoPC using intestinal epithelial Caco-2, known to be viable over a wide range of oxygen environments. In Fig. 5a, we follow OoC oxygen tension over the critical cell attachment phase. With NanoPC, readings remain stable around 15 kPa. This equilibrium is on the lower end of Zeitouni et al.’s report from “normoxic” well-plate Caco-2 cultures (16∼18.6 kPa; independent of standard or gas-permeable plates). While the calibration error noted earlier again plays a role here, the primary factor for our lower equilibrium likely lies in our higher cell density (relative to media volume). In stark contrast to NanoPC devices, the control PC devices become rapidly hypoxic in ∼30 minutes, dropping below even the Caco-2-relevant gastrointestinal-physioxic regime and remaining there throughout the experiment. We do note some outliers which we attribute to air bubbles or device leakage (high oxygen with control PC), and to contaminated membranes (e.g., glue residue clogging the pores; low oxygen with NanoPC).

**Figure 5.**
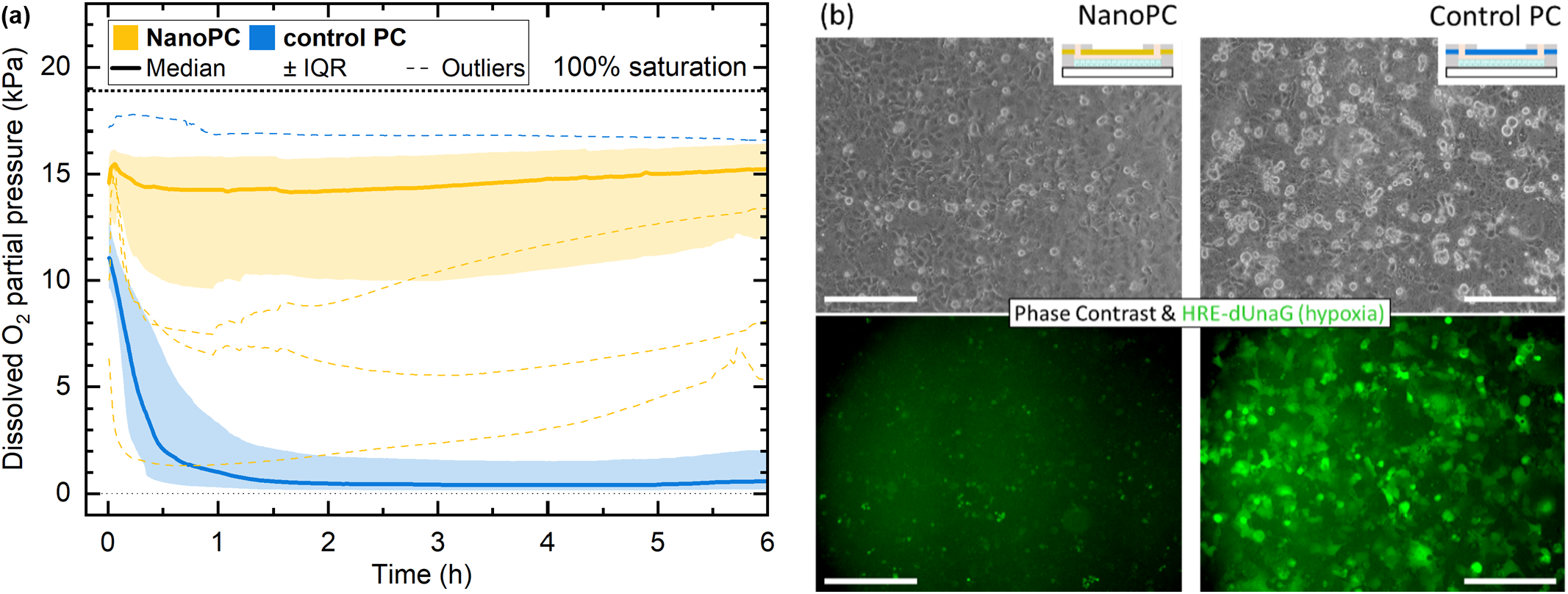
NanoPC-OoC evaluation upon seeding with Caco-2 cells. (a) Oxygen tension recordings from respective tape OoC, illustrating stable conditions with NanoPC-integrated ones, in clear contrast to rapid oxygen depletion with control PC. The solid lines represent the median of bootstrap means across 23 OoC each, including outliers (dotted lines; as per Tukey fences), and shaded regions indicating corresponding IQR. (b) Representative phase contrast and fluorescence micrographs from NanoPC-(left) and control PC (right)-based OoC after ∼26 h. Hypoxia response element fluorescence is markedly upregulated with NanoPC. Scale bars: 250 µm. Additional images in Fig. S5.

To confirm that these differences in oxygen tension impacted cellular function, we rely on a fluorescent reporter for HIF-1. In Fig. 5b, we can observe a distinct increase in fluorescence, indicating enhanced HIF-1 transcriptional activity as reported by the HRE-dUnaG system that would reflect oxygen-dependent stabilization of HIF-1α under hypoxic conditions, i.e., increased expression and/or reduced degradation of HIF-1α. The signal was marginal at 6 h (not shown) but became substantial overnight. This is consistent with the described delay of maturation and accumulation of this oxygen reporter element: In the original report for hypoxic Chinese hamster ovary (CHO) cells, a sigmoidal inflection point in signal intensity can be inferred around 10 h, with signals continuing to increase thereafter.[30] The underlying increase in HIF-1α occurs on a much shorter timescale; Zeitouni et al. reported clear upregulation of HIF-1α in hypoxic Caco-2 already within the first hours (though they did not determine when peak expression is reached).[49] The experiment confirms that the oxygen depletion in PC controls drives significant HIF-1 pathway engagement.

## Conclusion

We set out to address the continued lack of gas-permeable materials suitable for PDMS-free organ-on-a-chip fabrication — a gap that persists even as thermoplastic microfluidics have otherwise gained considerable traction, and a need that persists even as incubator-independent oxygenation management is emerging. To that end, we evaluated commercially available track-etched nanoporous polycarbonate (termed NanoPC herein), and found it to be an order of magnitude more oxygen permeable than PDMS. Four orders of magnitude more oxygen permeable than native PC, NanoPC nonetheless inherits PC’s mechanical properties, straightforward fabrication and integration capabilities, and low cost (more expensive than native PC, but still only ∼0.1 €/cm^2^). These compare favorably with alternatives, whether commercially-available silicone-free gas exchange membranes like porous fluoropolymers, or microfluidic literature-considered ones such as microporous silicon nitride or anodized aluminum oxide. We demonstrated NanoPC device integration using tape- or OSTEmer-lamination process flows, providing leak-free operation well above OoC-relevant pressures. Using Caco-2 intestinal epithelial cells, we validated the OoC application by monitoring oxygen levels during the cell attachment phase: NanoPC devices maintained a stable oxygen equilibrium over 6 h, whereas nonporous controls became severely and uncontrolledly hypoxic within 30 minutes. We moreover confirmed the cell-level impact as indicated by a hypoxia reporter element, where a clear hypoxia-inducible factor-1 response developed over 24 h of culture in the controls.

We identify and acknowledge a key limitation in that the nanoporosity increased not only oxygen permeability, but also water vapor permeance. Although the increase is less pronounced than for oxygen, and although thickness-normalized water vapor permeability is comparable to PDMS, we found this to be sufficient to impact cell viability. This demanded more careful humidity management in the immediate device vicinity than is typical. Mitigation approaches such as additional liquid reservoirs in close proximity (ours), or a limited-gas-volume buffer chamber (the presumed mitigating factor for Bunge et al.’s nanoporous membrane [27]) may not be practical in all contexts. One option lies in choosing NanoPC with an up to ∼10-fold lower porosity than our 1.18%, which would still place it on par with PDMS for oxygen permeability and be sufficient for many applications. Another opportunity we intend to pursue lies in spin-coating NanoPC with PDMS, wherein NanoPC would facilitate microfluidic integration and prevent direct PDMS-media contact (and thus small-molecule sorption), with PDMS contributing an easily thickness-tunable water vapor barrier. We moreover recognize that biological characterization with only Caco-2 cultures (i.e., a robust cancer cell line) and only for cell viability and hypoxia response is far from exhaustive. Yet, given the widespread use of PC in cell culture, we see little reason to expect biocompatibility complications even with iPSC-derived or primary cultures in future studies with suitable humidity management.

Ultimately, NanoPC holds great promise for the OoC field as it decouples gas permeability from PDMS limitations (small-molecule sorption, hydrophobicity, non-silicon-based material integration), and thus removes one of the two major reasons for continued reliance on silicone/PDMS. With thermoplastic elastomers addressing the other (i.e., elasticity; e.g., SEBS, PMP, PU), integration of both properties into the same PDMS-free device also becomes feasible. NanoPC as an external microfluidic material can provide stable incubator-controlled oxygenation with all-thermoplastic OoC, addressing also the pre-perfusion window where such devices otherwise leave cells vulnerable to uncontrolled hypoxia. At the same time, it can find application as an OoC-internal oxygen exchange membrane where regional or per-compartment oxygen control, independent of an incubator, is desired. Given NanoPC’s affordability and its favorable handling and integration characteristics – both in the lab, and at industrial scale – we see it not as a niche membrane material, but as a missing piece in the thermoplastic OoC toolkit that will help make controlled oxygen availability the norm rather than the exception.

## Acknowledgements

We greatly appreciate the experimental assistance and discussions regarding baseline oxygen membrane permeability with Stefanie Fuchs and Torsten Mayr (TU Graz & Pyroscience AT). We acknowledge funding by the European Union under ERC StG #101116523 “CHIPzophrenia.” Views and opinions expressed are, however, those of the author(s) only and do not necessarily reflect those of the European Union or the European Research Council. Neither the European Union nor the granting authority can be held responsible for them. We further acknowledge zukunft.niedersachsen, a funding program of the Lower Saxony Ministry of Science and Culture and the Volkswagen Foundation, within the MatrixSense project framework.

## Author Contributions

F.B. and J.B. contributed equally this work.

Conceptualization: T.E.W. (lead), J.B. & F.B. (supporting)

Data Curation & Formal Analysis & Methodology: F.B. & J.B. & T.E.W. (equal), G.K. & I.S. (supporting) Investigation & Validation: F.B. & J.B. (equal), G.K. & I.S. (supporting)

Resources: J.B. (lead), F.B., M.H. (supporting) Visualization: J.B. & F.B. & T.E.W. (equal)

Writing – original draft: T.E.W. & F.B. (equal), J.B. (supporting)

Writing – review & editing: T.E.W. & J.B. (equal), F.B. & M.H. (supporting)

Funding Acquisition & Project Administration: T.E.W. (lead), A.L. (supporting)

## References

[1] A. Carreau, B. E. Hafny-Rahbi, A. Matejuk, C. Grillon, and C. Kieda, “Why is the partial oxygen pressure of human tissues a crucial parameter? Small molecules and hypoxia,” J. Cell. Mol. Med., vol. 15, no. 6, pp. 1239–1253, 2011, doi: 10.1111/j.1582-4934.2011.01258.x.

[2] E. Ortiz-Prado, J. F. Dunn, J. Vasconez, D. Castillo, and G. Viscor, “Partial pressure of oxygen in the human body: a general review,” Am. J. Blood Res., vol. 9, no. 1, pp. 1–14, Feb. 2019.

[3] A. Krogh, “The supply of oxygen to the tissues and the regulation of the capillary circulation,” J. Physiol., vol. 52, no. 6, pp. 457–474, May 1919, doi: 10.1113/jphysiol.1919.sp001844.

[4] Y. Della Rocca et al., “Hypoxia: molecular pathophysiological mechanisms in human diseases,” J. Physiol. Biochem., vol. 78, no. 4, pp. 739–752, Nov. 2022, doi: 10.1007/s13105-022-00912-6.

[5] G. L. Semenza, “Regulation of Mammalian O2 Homeostasis by Hypoxia-Inducible Factor 1,” Annu. Rev. Cell Dev. Biol., vol. 15, no. Volume 15, 1999, pp. 551–578, Nov. 1999, doi: 10.1146/annurev.cellbio.15.1.551.

[6] V. Palacio-Castañeda, N. Velthuijs, S. Le Gac, and W. P. R. Verdurmen, “Oxygen control: the often overlooked but essential piece to create better in vitro systems,” Lab. Chip, vol. 22, no. 6, pp. 1068–1092, 2022, doi: 10.1039/D1LC00603G.

[7] J. A. Stuart et al., “How Supraphysiological Oxygen Levels in Standard Cell Culture Affect Oxygen-Consuming Reactions,” Oxid. Med. Cell. Longev., vol. 2018, no. 1, p. 8238459, Jan. 2018, doi: 10.1155/2018/8238459.

[8] T. L. Place, F. E. Domann, and A. J. Case, “Limitations of oxygen delivery to cells in culture: An underappreciated problem in basic and translational research,” Free Radic. Biol. Med., vol. 113, pp. 311–322, Dec. 2017, doi: 10.1016/j.freeradbiomed.2017.10.003.

[9] E. G. B. M. Bossink, L. I. Segerink, and M. Odijk, “Organ-on-Chip Technology for Aerobic Intestinal Host – Anaerobic Microbiota Research,” Organs---Chip, vol. 4, p. 100013, Dec. 2022, doi: 10.1016/j.ooc.2021.100013.

[10] P. E. Oomen, M. D. Skolimowski, and E. Verpoorte, “Implementing oxygen control in chip-based cell and tissue culture systems,” Lab. Chip, vol. 16, no. 18, pp. 3394–3414, 2016, doi: 10.1039/C6LC00772D.

[11] J. Santiago, J. Kreutzer, E. Bossink, P. Kallio, and J. le Feber, “Oxygen gradient generator to improve in vitro modeling of ischemic stroke,” Front. Neurosci., vol. 17, Mar. 2023, doi: 10.3389/fnins.2023.1110083.

[12] S. Fuchs, S. Johansson, A. Ø. Tjell, G. Werr, T. Mayr, and M. Tenje, “In-Line Analysis of Organ-on-Chip Systems with Sensors: Integration, Fabrication, Challenges, and Potential,” ACS Biomater. Sci. Eng., vol. 7, no. 7, pp. 2926–2948, Jul. 2021, doi: 10.1021/acsbiomaterials.0c01110.

[13] M. Azimzadeh, P. Khashayar, M. Amereh, N. Tasnim, M. Hoorfar, and M. Akbari, “Microfluidic-Based Oxygen (O2) Sensors for On-Chip Monitoring of Cell, Tissue and Organ Metabolism,” Biosensors, vol. 12, no. 1, p. 6, Jan. 2022, doi: 10.3390/bios12010006.A.L

[14] N. Jiang et al., “A closed-loop modular multiorgan-on-chips platform for self-sustaining and tightly controlled oxygenation,” Proc. Natl. Acad. Sci., vol. 121, no. 47, p. e2413684121, Nov. 2024, doi: 10.1073/pnas.2413684121.

[15] Z. Izadifar et al., “Organ chips with integrated multifunctional sensors enable continuous metabolic monitoring at controlled oxygen levels,” Biosens. Bioelectron., vol. 265, p. 116683, Dec. 2024, doi: 10.1016/j.bios.2024.116683.

[16] E. Zamir et al., “Dynamics and segregation of cell–matrix adhesions in cultured fibroblasts,” Nat. Cell Biol., vol. 2, no. 4, pp. 191–196, Apr. 2000, doi: 10.1038/35008607.

[17] S. Schlie, M. Gruene, H. Dittmar, and B. N. Chichkov, “Dynamics of Cell Attachment: Adhesion Time and Force,” Tissue Eng. Part C Methods, vol. 18, no. 9, pp. 688–696, Sep. 2012, doi: 10.1089/ten.tec.2011.0635.

[18] T. E. Winkler, J. S. Cognetti, B. M. Maoz, and A. Herland, “Materials for fabricating Micro-physiological Systems,” in Body-on-a-Chip, A. Atala and Y. Zhang, Eds., New York: Elsevier Academic Press, 2025.

[19] T. E. Winkler and A. Herland, “Sorption of Neuropsychopharmaca in Microfluidic Materials for In Vitro Studies,” ACS Appl. Mater. Interfaces, vol. 13, no. 38, pp. 45161–45174, Sep. 2021, doi: 10.1021/acsami.1c07639.

[20] J. Grant, A. Özkan, C. Oh, G. Mahajan, R. Prantil-Baun, and D. E. Ingber, “Simulating drug concentrations in PDMS microfluidic organ chips,” Lab. Chip, vol. 21, no. 18, pp. 3509–3519, Sep. 2021, doi: 10.1039/D1LC00348H.

[21] I. Ramos et al., “PDMS surface wettability modification and its applications: A systematic review,” J. Mol. Liq., vol. 434, p. 127978, Sep. 2025, doi: 10.1016/j.molliq.2025.127978.

[22] A. Gokaltun, M. L. Yarmush, A. Asatekin, and O. B. Usta, “Recent advances in nonbiofouling PDMS surface modification strategies applicable to microfluidic technology,” Technology, vol. 5, no. 1, pp. 1–12, Mar. 2017, doi: 10.1142/S2339547817300013.

[23] S. Seo and T. Kim, “Gas transport mechanisms through gas-permeable membranes in microfluidics: A perspective,” Biomicrofluidics, vol. 17, no. 6, p. 061301, Nov. 2023, doi: 10.1063/5.0169555.

[24] T. Hizawa, A. Takano, P. Parthiban, P. S. Doyle, E. Iwase, and M. Hashimoto, “Rapid prototyping of fluoropolymer microchannels by xurography for improved solvent resistance,” Biomicrofluidics, vol. 12, no. 6, p. 064105, Dec. 2018, doi: 10.1063/1.5051666.

[25] K. Giri and C.-W. Tsao, “Recent Advances in Thermoplastic Microfluidic Bonding,” Micromachines, vol. 13, no. 3, p. 486, Mar. 2022, doi: 10.3390/mi13030486.

[26] N. Arumugasaamy, J. Navarro, J. Kent Leach, P. C. W. Kim, and J. P. Fisher, “In Vitro Models for Studying Transport Across Epithelial Tissue Barriers,” Ann. Biomed. Eng., vol. 47, no. 1, pp. 1–21, Jan. 2019, doi: 10.1007/s10439-018-02124-w.

[27] F. Bunge, S. van den Driesche, and M. J. Vellekoop, “PDMS-free microfluidic cell culture with integrated gas supply through a porous membrane of anodized aluminum oxide,” Biomed. Microdevices, vol. 20, no. 4, p. 98, Nov. 2018, doi: 10.1007/s10544-018-0343-z.

[28] N. Imtiaz, W. A. Stoddard, A. Ghazy, and S. W. Day, “Oxygen transport in nanoporous SiN membrane compared to PDMS and polypropylene for microfluidic ECMO,” Biomed. Microdevices, vol. 27, no. 2, p. 22, May 2025, doi: 10.1007/s10544-025-00750-5.

[29] W.-I. Wu et al., “Lung assist device: development of microfluidic oxygenators for preterm infants with respiratory failure,” Lab. Chip, vol. 13, no. 13, pp. 2641–2650, Jun. 2013, doi: 10.1039/C3LC41417E.

[30] R. Erapaneedi, V. V. Belousov, M. Schäfers, and F. Kiefer, “A novel family of fluorescent hypoxia sensors reveal strong heterogeneity in tumor hypoxia at the cellular level,” EMBO J., vol. 35, no. 1, pp. 102–113, Jan. 2016, doi: 10.15252/embj.201592775.

[31] C. Schmitz et al., “Live reporting for hypoxia: Hypoxia sensor–modified mesenchymal stem cells as in vitro reporters,” Biotechnol. Bioeng., vol. 117, no. 11, pp. 3265–3276, 2020, doi: 10.1002/bit.27503.

[32] T. E. Winkler, M. Feil, E. F. G. J. Stronkman, I. Matthiesen, and A. Herland, “Low-cost microphysiological systems: feasibility study of a tape-based barrier-on-chip for small intestine modeling,” Lab. Chip, vol. 20, no. 7, pp. 1212–1226, Mar. 2020, doi: 10.1039/D0LC00009D.

[33] I. Matthiesen, D. Voulgaris, P. Nikolakopoulou, T. E. Winkler, and A. Herland, “Continuous Monitoring Reveals Protective Effects of N-Acetylcysteine Amide on an Isogenic Microphysiological Model of the Neurovascular Unit,” Small, vol. 17, no. 32, p. 2101785, Aug. 2021, doi: 10.1002/smll.202101785.

[34] K. Corral-Nájera, G. Chauhan, S. O. Serna-Saldívar, S. O. Martínez-Chapa, and M. M. Aeinehvand, “Polymeric and biological membranes for organ-on-a-chip devices,” Microsyst. Nanoeng., vol. 9, no. 1, p. 107, Aug. 2023, doi: 10.1038/s41378-023-00579-z.

[35] P. Apel, “Track etching technique in membrane technology,” Radiat. Meas., vol. 34, no. 1–6, pp. 559–566, Jun. 2001, doi: 10.1016/S1350-4487(01)00228-1.

[36] J. E. Mark, Ed., Polymer Data Handbook: Second Edition. Oxford University Press, 2009. doi: 10.1093/oso/9780195181012.001.0001.

[37] F. J. Norton, “Gas permeation through lexan polycarbonate resin,” J. Appl. Polym. Sci., vol. 7, no. 5, pp. 1649–1659, Nov. 1963, doi: 10.1002/app.1963.070070507.

[38] L. Sønstevold, M. Czerkies, E. Escobedo-Cousin, S. Blonski, and E. Vereshchagina, “Application of Polymethylpentene, an Oxygen Permeable Thermoplastic, for Long-Term on-a-Chip Cell Culture and Organ-on-a-Chip Devices,” Micromachines, vol. 14, no. 3, p. 532, Feb. 2023, doi: 10.3390/mi14030532.

[39] S. Zips et al., “Biocompatible, Flexible, and Oxygen-Permeable Silicone-Hydrogel Material for Stereolithographic Printing of Microfluidic Lab-On-A-Chip and Cell-Culture Devices,” ACS Appl. Polym. Mater., vol. 3, no. 1, pp. 243–258, Jan. 2021, doi: 10.1021/acsapm.0c01071.

[40] I. Blume, P. J. F. Schwering, M. H. V. Mulder, and C. A. Smolders, “Vapour sorption and permeation properties of poly (dimethylsiloxane) films,” J. Membr. Sci., vol. 61, pp. 85–97, Jan. 1991, doi: 10.1016/0376-7388(91)80008-T.

[41] K. Domansky et al., “SEBS elastomers for fabrication of microfluidic devices with reduced drug absorption by injection molding and extrusion,” Microfluid. Nanofluidics, vol. 21, no. 6, p. 107, Jun. 2017, doi: 10.1007/s10404-017-1941-4.

[42] E. Tech, “Flexdym for Fast & Easy Microfluidic Device,” EDEN TECH. Accessed: Feb. 10, 2026. [Online]. Available: https://eden-microfluidics.com/news-events/flexdym-for-fast-easy-microfluidic-device-fabrication/

[43] Y. Wang, M. Gupta, and D. A. Schiraldi, “Oxygen permeability in thermoplastic polyurethanes,” J. Polym. Sci. Part B Polym. Phys., vol. 50, no. 10, pp. 681–693, 2012, doi: 10.1002/polb.23053.

[44] I. Coelho, R. F. Pires, S. B. Gonçalves, V. D. B. Bonifácio, and M. Faria, “Gas Permeability and Mechanical Properties of Polyurethane-Based Membranes for Blood Oxygenators,” Membranes, vol. 12, no. 9, p. 826, Sep. 2022, doi: 10.3390/membranes12090826.

[45] Velianti, S. B. Park, and P. W. Heo, “The enhancement of oxygen separation from the air and water using poly(vinylidene fluoride) membrane modified with superparamagnetic particles,” J. Membr. Sci., vol. 466, pp. 274–280, Sep. 2014, doi: 10.1016/j.memsci.2014.04.043.

[46] A. Park et al., “Blood Oxygenation Using Fluoropolymer-Based Artificial Lung Membranes,” ACS Biomater. Sci. Eng., vol. 6, no. 11, pp. 6424–6434, Nov. 2020, doi: 10.1021/acsbiomaterials.0c01251.

[47] D. Sticker, R. Geczy, U. O. Häfeli, and J. P. Kutter, “Thiol–Ene Based Polymers as Versatile Materials for Microfluidic Devices for Life Sciences Applications,” ACS Appl. Mater. Interfaces, vol. 12, no. 9, pp. 10080–10095, Mar. 2020, doi: 10.1021/acsami.9b22050.

[48] A. M. Kemas et al., “Compound Absorption in Polymer Devices Impairs the Translatability of Preclinical Safety Assessments,” Adv. Healthc. Mater., vol. 13, no. 11, p. 2303561, 2024, doi: 10.1002/adhm.202303561.

[49] N. E. Zeitouni, J. Fandrey, H. Y. Naim, and M. von Köckritz-Blickwede, “Measuring oxygen levels in Caco-2 cultures,” Hypoxia, vol. 3, pp. 53–66, Oct. 2015, doi: 10.2147/HP.S85625.

