## Supplementary Information for "Overcoming oxygen impermeability in PDMS-free organ-on-a-chip microfluidics with nanoporous thermoplastic"

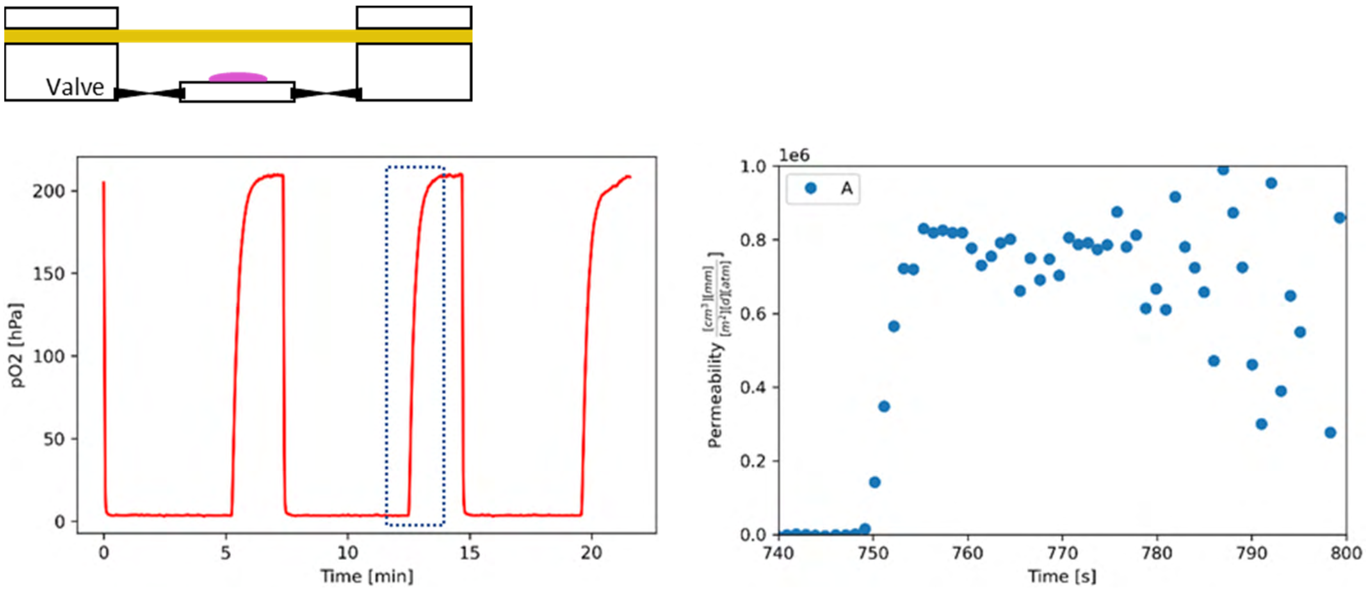


***Figure S1.*** *Top: oxygen permeability test cell, with the oxygen sensor spot (purple) recording partial oxygen pressure $p_{O2}$ over time as it is alternately flushed with nitrogen and allowed to re-equilibrate with room atmosphere. Bottom left: Exemplary raw measurement data trace including three experimental cycles from a single sample. From this, we calculated Permeability as* $P_{O_2}(t) = \frac{V\,d}{A}\frac{\frac{d}{dt}p_{O_2}(t)}{p_{atm}-p_{O_2}(t)}$, *where V is the measurement cell volume, A the area of the membrane sample, and $p_{atm}$ the partial oxygen pressure measured in room temperature air (i.e., upper-level sensor calibration for this experiment). Bottom right: Thus-derived oxygen permeability corresponding to the area highlighted with the blue dotted box. The measurement does not give stable results for more than 20–30 s as the oxygen level inside the setup is rapidly saturating. Averaging over the stable measurement range, and across replicates, yields our reported $P_{O2}$.*

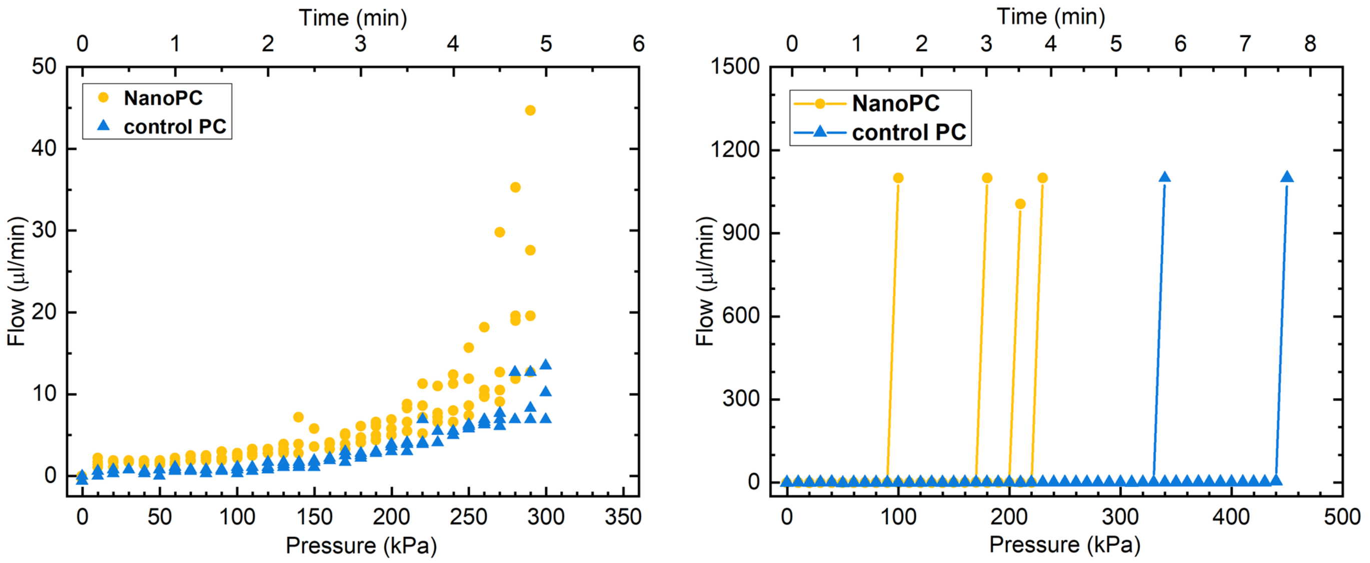


***Figure S2.*** *Short-term/burst pressure testing. Given the limitations of the flow sensor at low flows, we defined "baseline" (i.e., constant, near-zero flow) and "failure" (i.e., increasing flow) regimes, and defined the failure point as the intersection point of the increasing-flow linear fit with the baseline mean plus 1.96 SD. For tape devices (left), the baseline was defined by pointwise expansion of an initial 10–50 kPa range until new points exceeded 1.96 SD, and failure interval for linear fitting was defined as the subsequent 16 readings (chosen based on* $R^2$ *across all conditions). The different failure mode with OSTEmer (right) necessitated a different approach, wherein the failure interval was simply defined as the last two readings.*

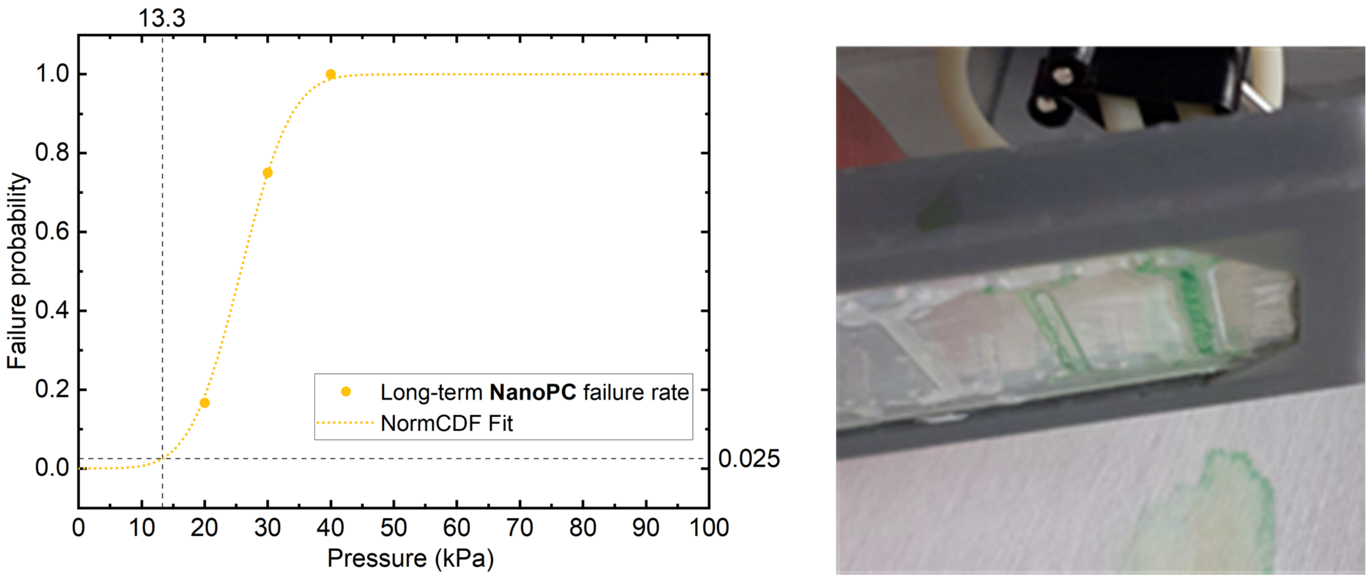


***Figure S3.*** *Long-term pressure testing. Left: Normal cumulative distribution function-fit of the long-term failure rates of NanoPC devices. The 2.5% (=1.96σ) failure point is indicated at 13.3 kPa. Right: Exemplary image of leaking device.*

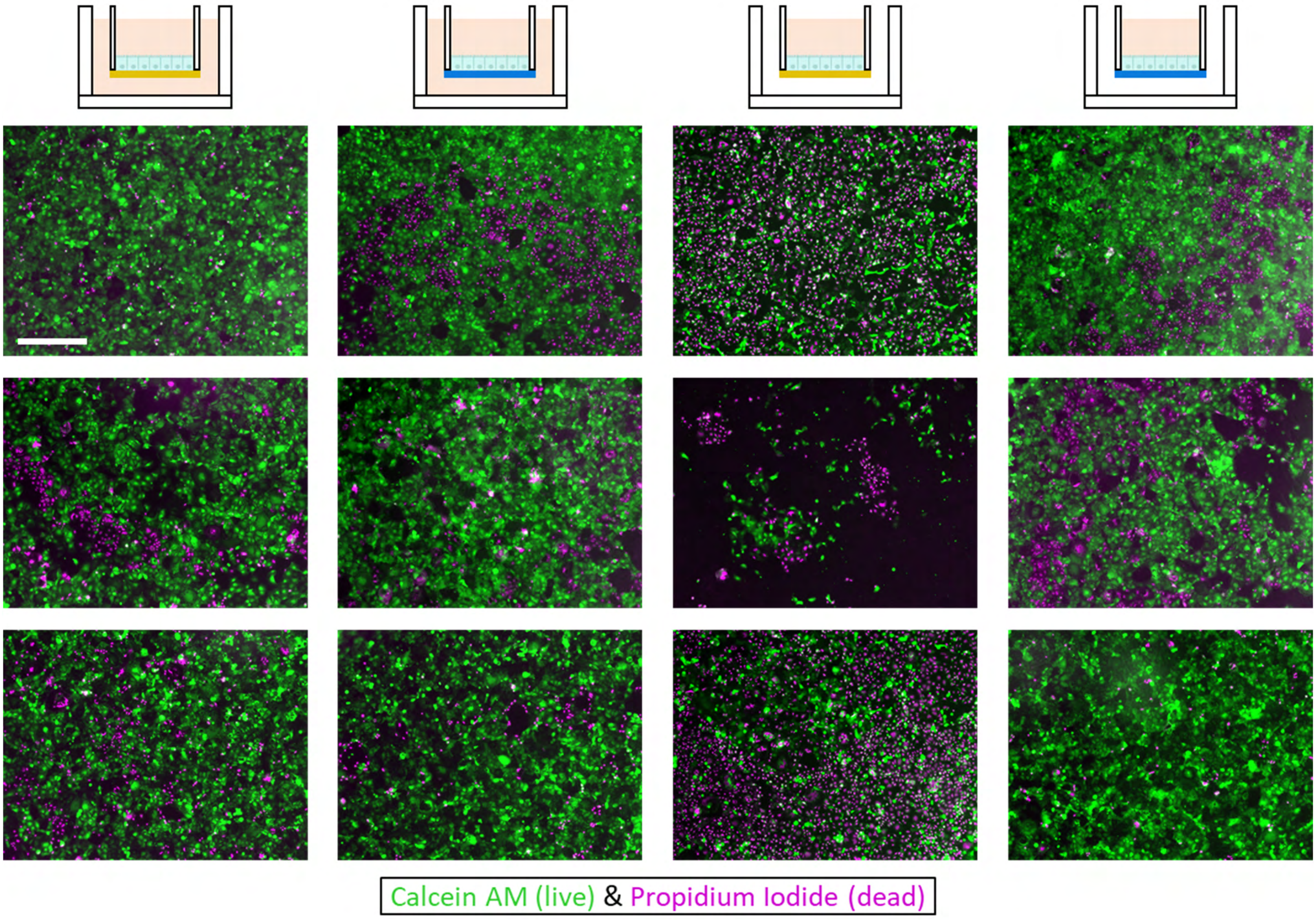


***Figure S4***. *Cytotoxicity evaluation of NanoPC (yellow; columns 1 and 3) as compared to PC controls (blue; columns 2 and 4) employed as cell culture substrates in Transwell-like geometry. Caco-2 cells are fluorescently stained with a membrane-permeable dye-linked esterase substrate (green; live cells), and a membrane-impermeable DNA-intercalating dye (magenta; dead cells). For each independent round of experiments (rows), we show one random number generator-selected micrograph per condition. Images reveal no inherent difference between NanoPC and control when immersed in liquid (left); with air underneath (right), NanoPC performs significantly worse, suggesting a potential water-evaporation effect. Scale bars: 500 µm.*

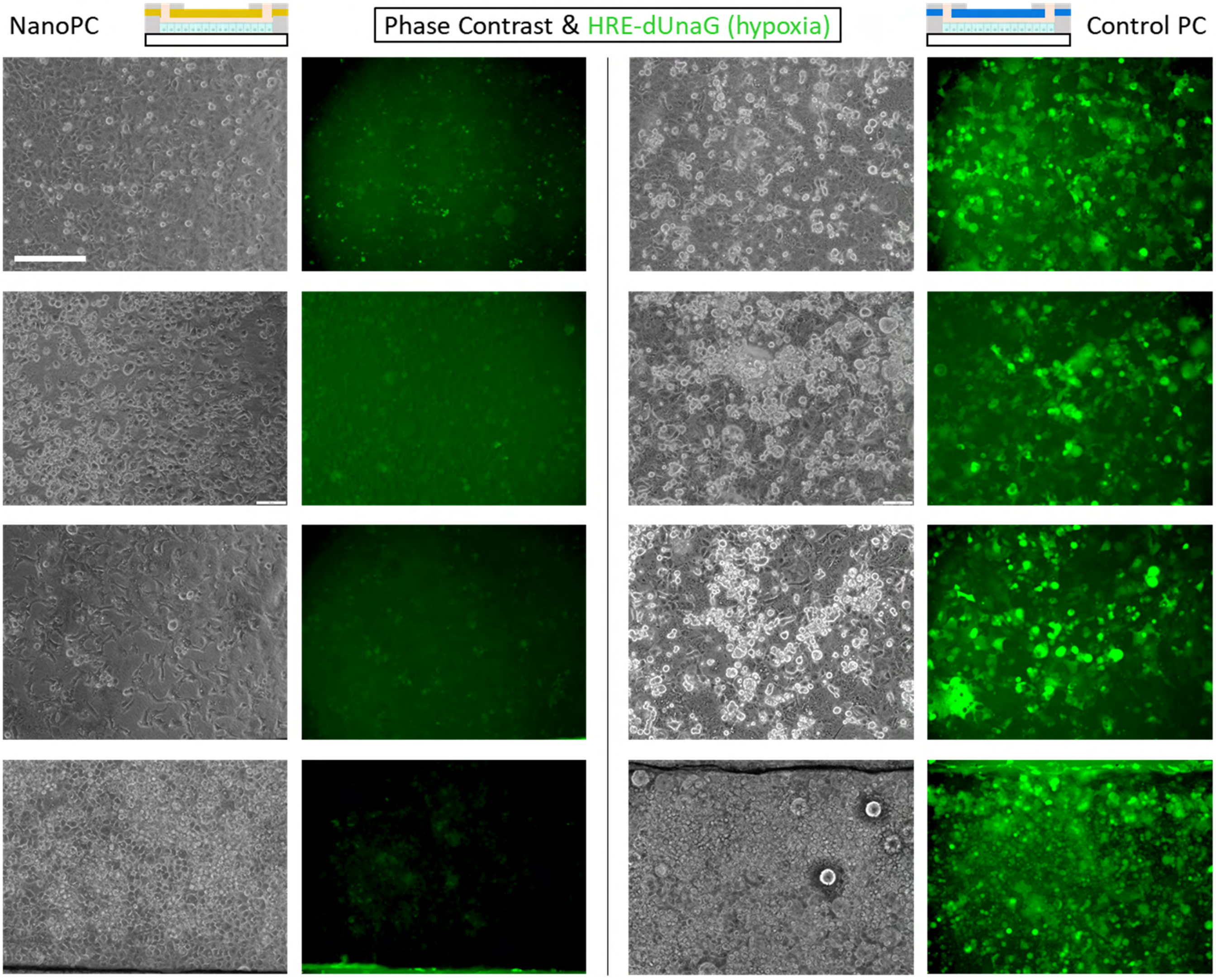


***Figure S5***. *Phase contrast and fluorescence micrographs from NanoPC- (left) and control PC (right)-based OoC after ~26 h. Engineered hypoxia response element fluorescence in the Caco-2 appears in green. For each independent round of experiment (rows), we show one exemplary image from around the middle (length-wise) of the microchannel, with the top row identical to Figure 5. Hypoxia response element fluorescence is markedly upregulated with NanoPC. Scale bars: 250 µm*

### Theoretical oxygen permeability

We calculate the expected oxygen permeability of our NanoPC membranes in terms of $P = \frac{D_{\text{eff}}}{R\,T}$, where $D_{\text{eff}}$ is the effective diffusion coefficient, $R$ the molar gas constant, and $T$ the temperature (for the purposes of Table 1, 23°C to match the initial experiment, and 37°C to match the ultimate application). For $D_{\text{eff}}$, we need to consider three factors: (1) the fundamental restriction inherent in the free area fraction or porosity $\varepsilon$ (since our pores are straight, no tortuosity correction, often similarly present for porous media, is needed); as well as (2) bulk gas diffusion, i.e., limited by collisions with other gas molecules; and/or (3) the restriction inherent in the confined space of each single pore. To determine the relevance of the latter two, we can calculate the Knudsen number $K = \lambda/d$ of 1.33, based on a mean free oxygen path length of $\lambda$=66 nm (at 23°C).[1] Being close to 1 (even across a range of temperatures), we thus need to consider both bulk diffusion and Knudsen diffusion as per the Bosanquet equation [2] i.e., $D_{\text{eff}} = \frac{\varepsilon}{\frac{1}{D_{O_2}}+\frac{1}{D_{\text{pore}}}}$, with $D_{O_2} = \left(\frac{T}{273.15\text{ K}}\right)^{1.81} 0.182\ \text{cm}^2/s$ the estimated bulk diffusivity of oxygen in air, [3] and $D_{\text{pore}} = \frac{d}{3}\sqrt{\frac{8\,R\,T}{\pi\,M_{O_2}}}$ the Knudsen diffusion in a confined pore,[4] wherein $M_{O2}$ is the molar mass of oxygen.

### Device fabrication: Pressure resilience testing

These microfluidic devices (Fig. 2a) were built on a commercial polycarbonate fluidic interface platform (Fluidic 343, Microfluidic ChipShop, Germany) with double-sided tape or OSTEmer.

Tape devices were constructed following the method outlined by Winkler et al. [5]. In brief, channels were cut out of double-sided medical grade polyester tape (9877, 3M) using a cutter plotter (M1, xTool). The channels are ~22 mm long and 1.5 mm wide. A channel height of ~200 µm was achieved by using 2 layers of tape. Two such channel layers were aligned and stuck to the fluidic connector plate. To close the system, either NanoPC or native PC was stuck to the bottom of the system. After assembling the tape and PC layers, the devices were clamped together between glass slides with paper clamps, and left in vacuum overnight, to ensure a good seal.

For the geometrically identical OSTEmer devices, the protocol described by Matthiesen et al. [6] was largely followed. In brief: A positive mold machined in aluminium and coated with parylene was used to cast a negative PDMS mold. OSTEmer 322 (Mercene AB) was reaction injection molded, with curing initiated by 365 nm light exposure to create two channel layers. After the first curing reaction, the channels were demolded, aligned, and sandwiched between the fluidic connector platform and either NanoPC or native PC. These parts were then clamped together and the second curing step was performed by baking at 120°C overnight.

### Device fabrication: Oxygen permeability testing

The tape-based chips were adapted for integration of oxygen sensor spots (Fig. 2b). Before cutting, a layer of tape was effectively rendered one-sided by sticking a 1 µm thick layer of Mylar C (DuPont) against one side. This step prevents the channel top and bottom from inadvertently bonding together. Then through-holes for inlets and a rectangular hole (2.7×2 mm$^2$) were cut with a cutter plotter. This layer was stuck to the same commercial connector plate as before. In the rectangular cutout, a manually cut piece of an oxygen sensor spot (Pyroscience, OXSP5) was glued against the connector plate with PDMS (Sylgard 184, 1:10, cured at 60°C for 1h). Channels (~22×1.5 mm$^2$) were cut out of single-layer double-sided tape and added to the plate, forming channels of ~100 µm high, with a sensor spot in a rectangular recess in the channel ceiling at the channel center. The system was again closed with either native PC or NanoPC.

### Device fabrication: Cell culture testing

For the devices for measuring gas exchange during cell culture (Fig. 2c), the tape-based process was adapted further, cutting three layers out of double-sided medical tape. For the upper layer, a single layer of tape was used. In this layer a singular channel of ~18×3 $mm^2$ was cut. For the middle layer, a single layer of tape (rendered one-sided as above) was used. From the middle layer, 4 through-holes and 1 central circular hole for the sensor spot were cut. The bottom layer was cut out of a double layer of tape. From this layer 2 channels of ~8×1 $mm^2$ were cut to connect the through-holes in the middle layer. A larger circular hole was cut from the middle. Whole sensor spots were glued with PDMS to a circular laser-cut native PC. This assembly was then aligned so that the sensor spot fit through the hole in the middle channel, while the carrier PC was stuck to the adhesive of the middle layer. The channels in the lower layers were sealed with native PC. Afterwards, 3D printed reservoirs (Printer: Form 4; Resin: High Temp; Formlabs) were attached to the outer through-holes of the middle layer.

### Well plate devices

To construct devices suitable for cell culture observation (Fig. 2d), equivalent-geometry channels (~18×3 $mm^2$) were cut out of a single layer of double-sided tape. One side was sealed with either NanoPC or native PC. 3D printed reservoirs were stuck on top using double-sided tape, and through-holes were manually cut. This assembly was then stuck to the bottom of a well of a standard 6-well cell culture plate (Sarstedt, Germany).

### Evaporation testing

PC, NanoPC or PDMS membranes were clamped over 1.9 ml volume 3D printed vessels (Printer: Form 4; Resin: White V5; Formlabs) filled with 1 ml water and sealed with an O-ring. Vessels without membrane served as control. The PDMS membranes were manufactured from Sylgard 184 in a 10:1 ratio by spin coating a 4-inch glass wafer at 550 rpm for 5 minutes and curing at 120°C for 3 hours. The vessels were weighed and left for 24 hours in a controlled climate at 22°C and 23-28% relative humidity, after which they were weighed again to determine the amount of evaporated water.

### Cell culture

Frozen stocks of human enterocytes (Caco-2 BBe1, or Caco-2 transduced with HRE-dUnaG [7][8]) were expanded and maintained at 37 °C / 5% CO2 according to supplier protocols. Media was prepared from DMEM (high glucose; Gibco 10569010) and 100 U/ml penicillin–streptomycin (Gibco 15140122).

For the cytotoxicity assay, we supplemented the media with 20% heat-inactivated fetal bovine serum (FBS; Gibco A3840002). In the remainder of our study, due to rising costs, we employed 10% FBS combined with 1× insulin-transferrin-selenium (ITS; Gibco 41400045). Media was prepared the day prior to use and placed in the incubator overnight to equilibrate. Pre-equilibration to 37 °C and to the “correct” partial pressures of dissolved gases reduces one potential source of bubble formation.

Devices in any cell experiment were disinfected with 70% ethanol, flushed with PBS, and coated with collagen I (Bio-Techne, USA) diluted in 20 mM acetic acid, either 0.1 g/L for 1–2 h (cytotoxicity; oxygen monitoring) or 0.5 g/L overnight (hypoxia observation), optimized empirically on a per-design basis. The coating solution was flushed out with PBS, followed by media, prior to seeding.

### Notes and references


[1] S. G. Jennings, “The mean free path in air,” *J. Aerosol Sci.*, vol. 19, no. 2, pp. 159–166, Apr. 1988, doi: 10.1016/0021-8502(88)90219-4.

[2] W. G. Pollard and R. D. Present, “On Gaseous Self-Diffusion in Long Capillary Tubes,” *Phys. Rev.*, vol. 73, no. 7, pp. 762–774, Apr. 1948, doi: 10.1103/PhysRev.73.762.

[3] W. J. Massman, “A review of the molecular diffusivities of H2O, CO2, CH4, CO, O3, SO2, NH3, N2O, NO, and NO2 in air, O2 and N2 near STP,” *Atmos. Environ.*, vol. 32, no. 6, pp. 1111–1127, Mar. 1998, doi: 10.1016/S1352-2310(97)00391-9.

[4] M. Knudsen, “Die Gesetze der Molekularströmung und der inneren Reibungsströmung der Gase durch Röhren,” *Ann. Phys.*, vol. 333, no. 1, pp. 75–130, Jan. 1909, doi: 10.1002/andp.19093330106.

[5] T. E. Winkler, M. Feil, E. F. G. J. Stronkman, I. Matthiesen, and A. Herland, “Low-cost microphysiological systems: feasibility study of a tape-based barrier-on-chip for small intestine modeling,” *Lab. Chip*, vol. 20, no. 7, pp. 1212–1226, Mar. 2020, doi: 10.1039/D0LC00009D.

[6] I. Matthiesen, D. Voulgaris, P. Nikolakopoulou, T. E. Winkler, and A. Herland, “Continuous Monitoring Reveals Protective Effects of N-Acetylcysteine Amide on an Isogenic Microphysiological Model of the Neurovascular Unit,” *Small*, vol. 17, no. 32, p. 2101785, Aug. 2021, doi: 10.1002/smll.202101785.

[7] R. Erapaneedi, V. V. Belousov, M. Schäfers, and F. Kiefer, “A novel family of fluorescent hypoxia sensors reveal strong heterogeneity in tumor hypoxia at the cellular level,” *EMBO J.*, vol. 35, no. 1, pp. 102–113, Jan. 2016, doi: 10.15252/embj.201592775.

[8] C. Schmitz *et al.*, “Live reporting for hypoxia: Hypoxia sensor–modified mesenchymal stem cells as in vitro reporters,” *Biotechnol. Bioeng.*, vol. 117, no. 11, pp. 3265–3276, 2020, doi: 10.1002/bit.27503.
